# CD73-positive cell spheroid transplantation attenuates colonic atrophy

**DOI:** 10.1101/2022.08.23.505052

**Authors:** Daisuke Hisamatsu, Natsumi Itakura, Yo Mabuchi, Rion Ozaki, Eriko Grace Suto, Yuna Naraoka, Akari Ikeda, Lisa Ito, Chihiro Akazawa

## Abstract

The incidence of inflammatory bowel diseases (IBD) is increasing worldwide. Mesenchymal stem/stromal cells (MSCs) have immunomodulatory functions and are a promising source for cell transplantation therapy for IBD. However, owing to their heterogeneous nature, their therapeutic efficacy in colitis is controversial and depends on the delivery route and form of transplanted cells. Cluster of differentiation (CD)73 is widely expressed in MSCs and used to obtain a homogeneous MSC population. Herein, we determined the optimal method for MSC transplantation using CD73^+^ cells in a colitis model. mRNA sequencing analysis showed that CD73^+^ cells exhibit downregulation of inflammatory genes and upregulation of extracellular matrix-related genes. Furthermore, three-dimensional CD73^+^ cell spheroids showed enhanced engraftment at the injured site through the enteral route, facilitated extracellular matrix remodeling, and downregulated inflammatory gene expression in fibroblasts, leading to attenuation of colonic atrophy. Therefore, the interaction between intestinal fibroblasts and exogenous MSCs via tissue remodeling is one mechanism that can be exploited for colitis prevention. Our results highlight that transplantation of homogeneous cell populations with well-characterized properties is beneficial for IBD treatment.

## Introduction

Inflammatory bowel diseases (IBD), including ulcerative colitis (UC) and Crohn’s disease, are intractable diseases whose incidence rate in recent years has increased up to 0.5% of the population in industrialized countries, especially in Western countries (Holmberg *et al*, 2017). Moreover, in recent years, more than 180,000 people in Asia have been reported to have UC due to increased westernization of lifestyle, including food habits (Okabayashi *et al*, 2020). IBD are caused by chronic inflammation because of activated immune cells; however, no fundamental therapy exists for the condition (Gonzalez-Rey *et al*, 2009). Given the immunosuppressive effects of mesenchymal stem/stromal cells (MSCs), stem cell transplantation therapy for inflammatory diseases, including IBD, has recently attracted attention (Wang *et al*, 2016). Although intravenous, intraperitoneal, and anal injections are known routes for delivery of MSCs in a mouse model of dextran sulfate sodium salt (DSS)-induced colitis, their therapeutic efficacy is controversial (Goncalves Fda *et al*, 2014; Wang *et al*, 2016). MSCs form a highly heterogeneous cell population that can be isolated from either tissues or cells that attach to cultured dishes (Grace Suto *et al*, 2015). Depending on the molecules expressed on MSCs, their ability for engraftment at the transplant site and immune modulation via secreted factors may differ. Moreover, recent findings have shown that spheroids derived from three-dimensional (3D) MSC cultures have enhanced immunomodulatory function, vascularization, and multipotency via alteration of their gene expression profiles in several disease models (Bartosh *et al*, 2010; Bhang *et al*, 2012; Niibe *et al*, 2020). Thus, the controversial therapeutic efficacy of the transplant route for the colitis model could be attributed to inappropriate selection of the delivery method for the appropriate cell form.

We previously demonstrated that cluster of differentiation (CD)73 molecules can be used to efficiently isolate cell populations with MSC characteristics from human and rodent adipose tissues (Suto *et al*, 2017; Suto *et al*, 2020). However, the specific characteristics of CD73 ^+^ cells remain unclear. Moreover, the appropriate route of transplantation in the colitis model and forms of transplanted cells, such as two-dimensional (2D) or 3D culture-derived spheroids, are unknown.

Tissue-resident fibroblasts are involved in intestinal homeostasis in the steady state and under inflammatory conditions, such as in the pathogenesis of IBD (McCarthy *et al*, 2020; Pasztoi & Ohnmacht, 2022). In the steady state, fibroblasts produce extracellular matrix (ECM)-related molecules, such as collagen I, collagen IV, fibronectin, and matrix metalloproteinases (MMPs), and maintain intestinal architecture through ECM remodeling (Bonnans *et al*, 2014). Furthermore, fibroblasts expressing specific genes play a pivotal role in maintaining intestinal stem cells via Wnt signaling (Degirmenci *et al*, 2018; Valenta *et al*, 2016). In contrast, single-cell analyses of patients with UC have shown that inflammation-associated fibroblasts are a heterogeneous population divided into several subsets that can be classified according to the expression patterns of Wnt and bone morphogenic protein (BMP) family members (Kinchen *et al*, 2018; Smillie *et al*, 2019). These cells act in a context-dependent manner to either promote pathogenesis or suppress inflammation. Thus, the role of fibroblasts during development and pathogenesis of the intestinal tract remains to be clarified. However, the interaction between tissue-resident fibroblasts and exogenous MSCs during the acute or regeneration phases after MSC transplantation and the function of transplanted MSCs in intestinal homeostasis is largely unknown.

In the present study, we hypothesized that a uniform population of CD73^+^ cells, rather than conventional heterogeneous MSCs, would have more predictive or controlled therapeutic efficacy when considering cell transplantation at the clinical stage. Our goal was to characterize CD73^+^ cells by comparing them to conventional heterogeneous MSCs based on transcriptome analysis and determining the appropriate transplantation route and cell form for CD73^+^ cells in a mouse model of DSS-induced colitis. To this end, we analyzed gene and protein expression associated with intestinal homeostasis in the interactions between exogenous CD73^+^ cells and fibroblasts. Our study provides insights into MSC transplantation therapy for IBD and shows whether CD73^+^ cells are effective in this regard.

## Results

### Intravenous injection of CD73^+^ cells attenuates tissue destruction in DSS-induced colitis

To investigate the efficacy of CD73^+^ cell transplantation in a mouse model of DSS-induced colitis, we initially isolated CD45^−^CD31^−^Ter119^−^CD73^+^ (CD73^+^) cells from the subcutaneous white adipose tissue of mice using fluorescence-activated cell sorting (FACS) and compared them to conventional heterogeneous cell populations (from now on referred to as cMSCs) attached to the culture dish (**Appendix Fig S1 and Fig 1A**). In the present study, given the relevance of therapeutic strategies at an early stage before the colitis worsens, the first intravenous injection was administered a day after administration of DSS in drinking water, and the second dose was administered on day 5 (**Fig 1A**). After 3 days of recovery with regular water following DSS exposure, colon length was measured. Although there was no significant difference between the cMSC-transplanted and phosphate-buffered saline (PBS)-treated groups, the CD73^+^ cell-transplanted group had significantly longer colon length than the PBS-treated and cMSC-transplanted groups (p = 0.01, and p = 0.02, respectively; **Fig 1B**). Based on the colitis-associated histological score (Erben *et al*, 2014; Ernst *et al*, 2015), we evaluated the colon mucosa after hematoxylin and eosin (H&E) staining using the following criteria: mucosal architecture (0–3), immune cell infiltration (0–3), muscle layer thickness (0–3), depletion of goblet cells (0–1), and crypt abscess (0–1). The overall score of the CD73^+^ cell-transplanted group was lower than those of the cMSC-transplanted and PBS-treated groups (mean scores: CD73^+^ 5.6, cMSC 8.0, and PBS 10.3; **Figs 1C and D**). Specifically, the scores for mucosal architecture and crypt abscess were significantly lower in the CD73^+^ cell-transplanted group than those in the PBS-treated group (p = 0.05 and p = 0.02, respectively). Altogether, these results suggest that intravenous injection of CD73^+^ cells has more regenerative potency than intravenous injection of cMSCs in DSS-induced colitis.

**Figure 1.**
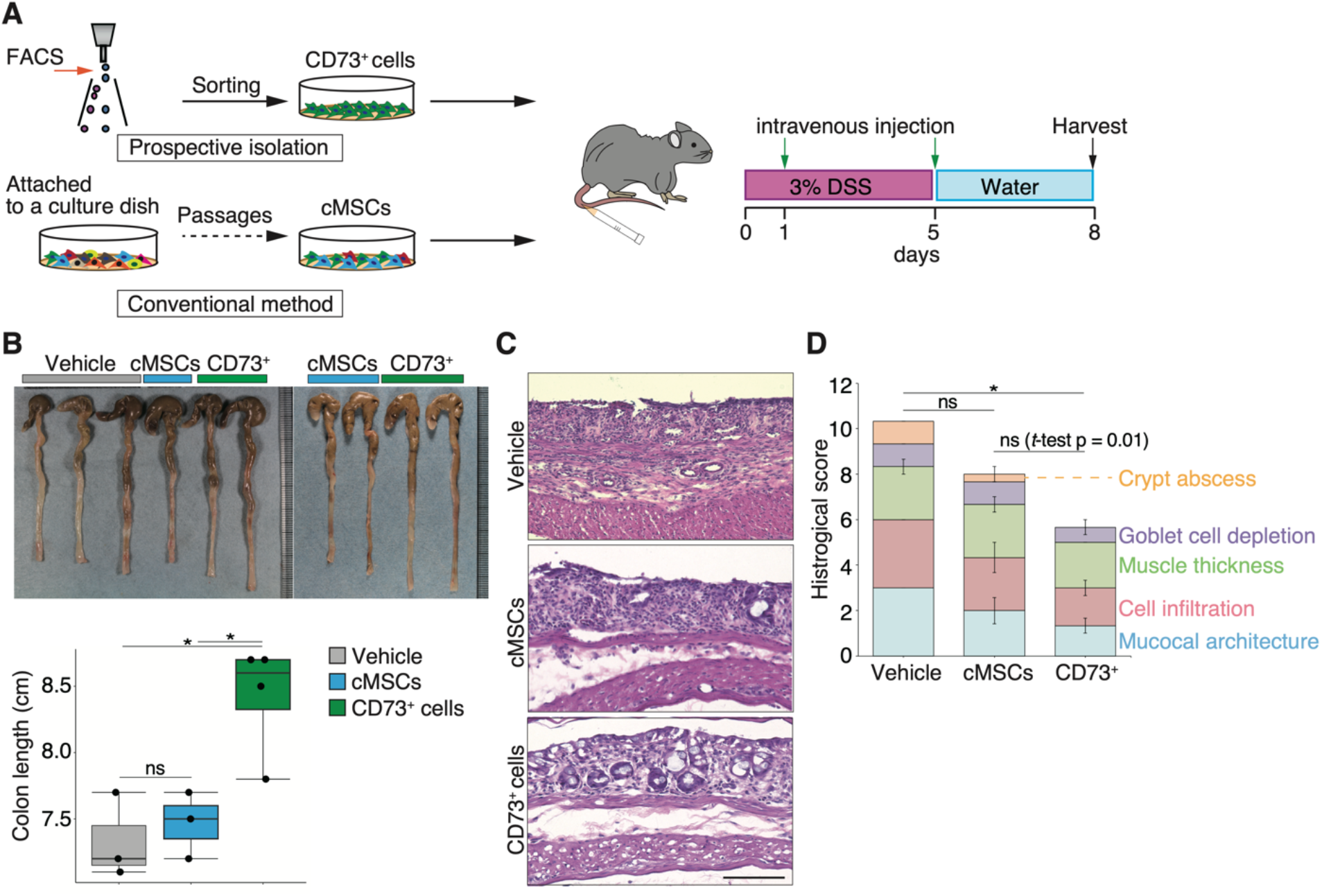
Intravenous injection of CD73^+^ cells attenuates destruction of intestinal structure in DSS-induced colitis. A Schematic diagram showing the isolation method and transplantation schedule of CD73^+^ cells and cMSCs. B Representative images of the colon after DSS-induced colitis and graph showing the colon length. Vehicle, PBS-treated group; cMSCs, cMSC-transplanted group; CD73^+^, CD73^+^ cell-transplanted group. C Representative images of H&E staining of the injured colonic tissue. Scale bar: 100 μm. D Graph showing histological score based on H&E staining of the colonic tissue. Each color represents the following criteria: mucosal architecture (0–3), immune cell infiltration (0–3), muscle layer thickness (0–3), depletion of goblet cells (0-1), and crypt abscess (0–1). Statistical significances were determined using Tukey’s test and Welch’s t-test. Data shown as mean ± standard error of mean (SEM). *p < 0.05; ***p < 0.005; ns, not significant.

### ECM-related genes are enriched in the CD73^+^ cell population

To characterize the highly regenerative efficacy of CD73^+^ cells, we performed a comparative transcriptome analysis between cMSCs and CD73^+^ cells isolated from human adipose tissue (**Appendix Fig S2**). We identified significantly differentially expressed genes that upregulated 882 and downregulated 1104 genes in CD73^+^ cells (log_2_ fold change > 1 or < −1, p < 0.05; **Fig 2A**). Gene ontology (GO) enrichment analysis revealed that the top 10 GO biological process terms, which include “GTP biosynthetic process,” “spliceosomal complex assembly,” “cell division,” “negative regulation of platelet-derived growth factor receptor signaling pathway,” and “mRNA transport,” were observed in the genes upregulated in CD73^+^ cells. In contrast, the terms for the downregulated genes were “immune response,” “innate immune response,” “adaptive immune response,” “inflammatory response,” and “neutrophil chemotaxis” (**Fig 2A**). We also found that *SMAD4*, fibronectin 1 *(FN1), S100A13, FBXO22*, and *SLC3A2*, known functional genes, were considerably upregulated in CD73^+^ cells (adjusted *P*-values < 0.05; **Fig 2A**). These genes are involved in integrin signaling and angiogenesis (Feral *et al*, 2007; Hayrabedyan *et al*, 2005; Schiller *et al*, 2004; Yang *et al*, 2020; Zheng *et al*, 2020), and the results are consistent with the results of gene set enrichment analysis (GSEA), in which the genes enriched in CD73^+^ cells were related to “regulation of collagen metabolic process,” “positive regulation of coagulation,” “extracellular matrix structural constituent,” and “transforming growth factor beta receptor binding” (**Fig 2B**). Furthermore, intravenously injected cells rarely engrafted into the colon (**Appendix Fig S3**), consistent with a previous study in which transplanted cells engrafted into the lungs (Hisamatsu *et al*, 2016). These results suggest that the regenerative efficacy of transplantation is due to the paracrine effect of secretory factors from engrafted cells at the transplant site. Therefore, we focused on the terms associated with paracrine signaling in our GO analysis results, and the “Wnt signaling pathway (GO:0016055)” was overrepresented in the genes upregulated in CD73^+^ cells (enrichment score: 1.22, p = 0.06).

**Figure 2.**
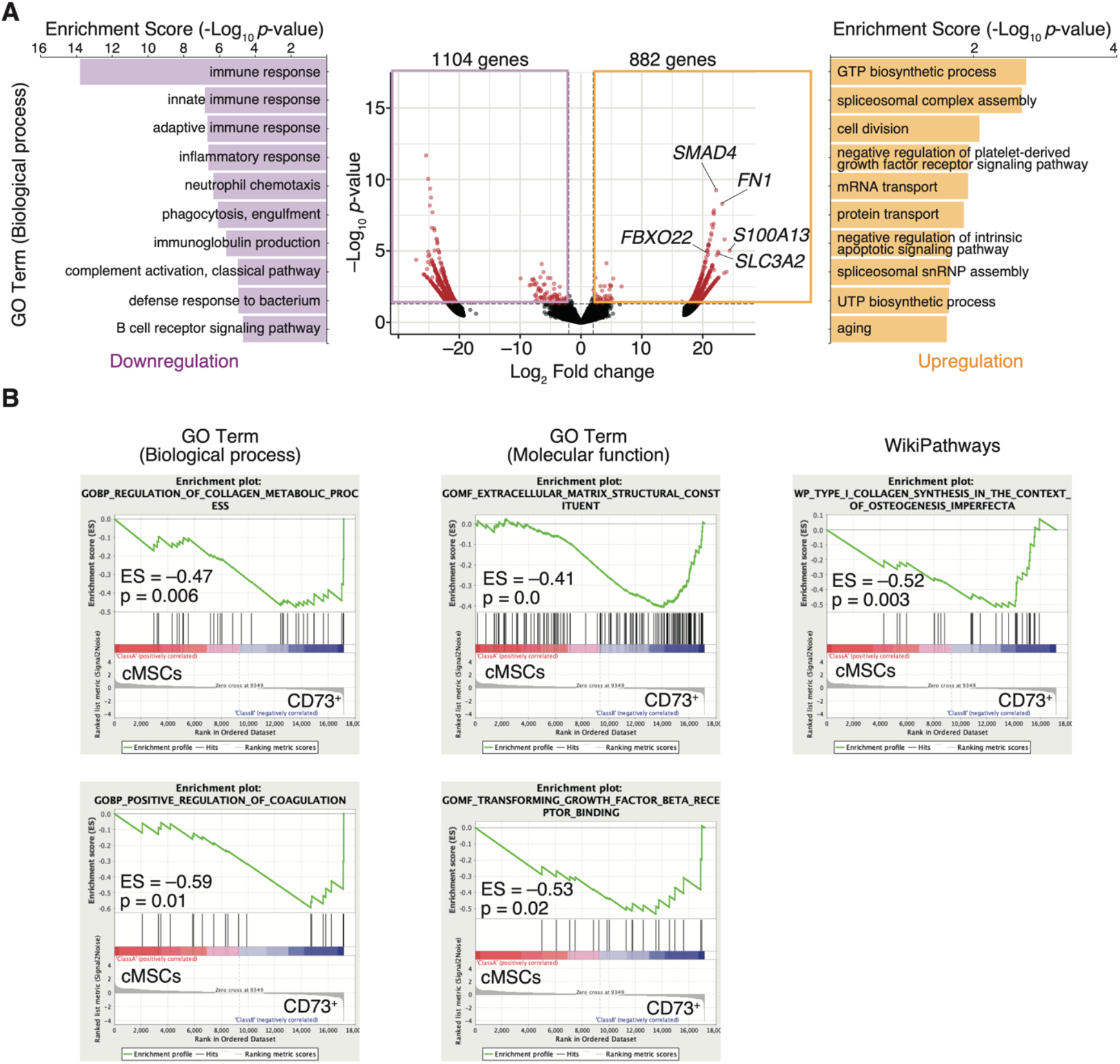
CD73^+^ cells exhibit upregulated gene expression related to ECM remodeling and downregulated gene expression related to immune response. A Volcano plot showing whole genes as differentially genes with a log2 fold change > 1 or < –1 (vertical dashed lines) and p < 0.05 (horizontal dashed line). Graphs showing the top 10 enrichment scores of GO biological process term by GO enrichment analysis. The left panel shows the downregulated genes, and the right panel shows the upregulated genes in CD73^+^ cells. B Gene set enrichment analysis of CD73^+^ cells and cMSCs. ES, enrichment score.

### Three-dimensional spheroids of CD73^+^ cells have more engraftment potential than two-dimensional cells in transanal transplantation

Considering that the genes related to cell adhesion, such as integrin signaling, were upregulated in CD73^+^ cells (**Fig 2B**), we hypothesized that CD73^+^ cells are suitable for engraftment to injured sites and attempted transanal transplantation, which is more accessible to the injured site in the distal colon than intravenous injection. Moreover, 3D culture-derived spheroid transplantation is associated with enhanced engraftment compared with 2D cell transplantation (Hu *et al*, 2020). In particular, 3D culture-derived intestinal organoids engraft in the injured colonic lumen after transanal administration (Fukuda *et al*, 2014). We first optimized the delivery method and compared a thin, flexible catheter with a 2.1 mm diameter to a stainless steel needle with a 1.9 mm diameter and found that the needle showed a higher survival rate than the catheter (**Appendix Figs S4A and B**). We further investigated differences in the engraftment rate of transplanted cells via the enteral route using a stainless steel needle, depending on the type of transplanted cells. On day 2 after transplantation of green fluorescent protein (GFP)-positive cells, the positivity rate was evaluated by FACS analysis of the distal colon tissue. We observed a significantly higher engraftment rate after spheroid transplantation than 2D cell transplantation (p = 0.01; **Appendix Figs S5A and B**).

### Transplantation of CD73^+^ cell spheroids onto the colonic lumen prevents mucosal atrophy

We found transanal transplantation of spheroids to be an effective delivery method for CD73^+^ cells, as indicated by the engraftment rate, but the therapeutic efficacy of this method on DSS-induced colitis is unknown. We prepared CD73^+^ cell-derived spheroids of 185.1 μm diameter (median) through two days of 3D culture (**Figs 3A and B**). To examine the effect of transplantation in the acute phase, we transferred these spheroids via the transanal route on days 0 and 5 of DSS administration and then measured the colon length on day 7 after DSS administration (**Fig 3C**). The colon length in the CD73^+^ spheroid-transplanted group was significantly higher than that in the PBS-and 5-aminosalicylic acid (5-ASA)-treated groups (5-ASA is used clinically for treating UC (Chapman *et al*, 2020)) on day 7 after DSS administration (p = 0.02 and p < 0.005, respectively; **Figs 3D**). Moreover, the proportion of endogenous GFP^−^CD45^−^CD140a^+^ fibroblasts in colonic tissue was significantly higher in the CD73^+^ spheroid-transplanted group than in the PBS- and 5-ASA treated groups on day 7 after DSS administration (p = 0.01 and p = 0.04, respectively; **Fig 3E**). These results suggest that CD73^+^ cell spheroid transplantation via the transanal route prevents the development of colitis atrophy during the acute phase.

**Figure 3.**
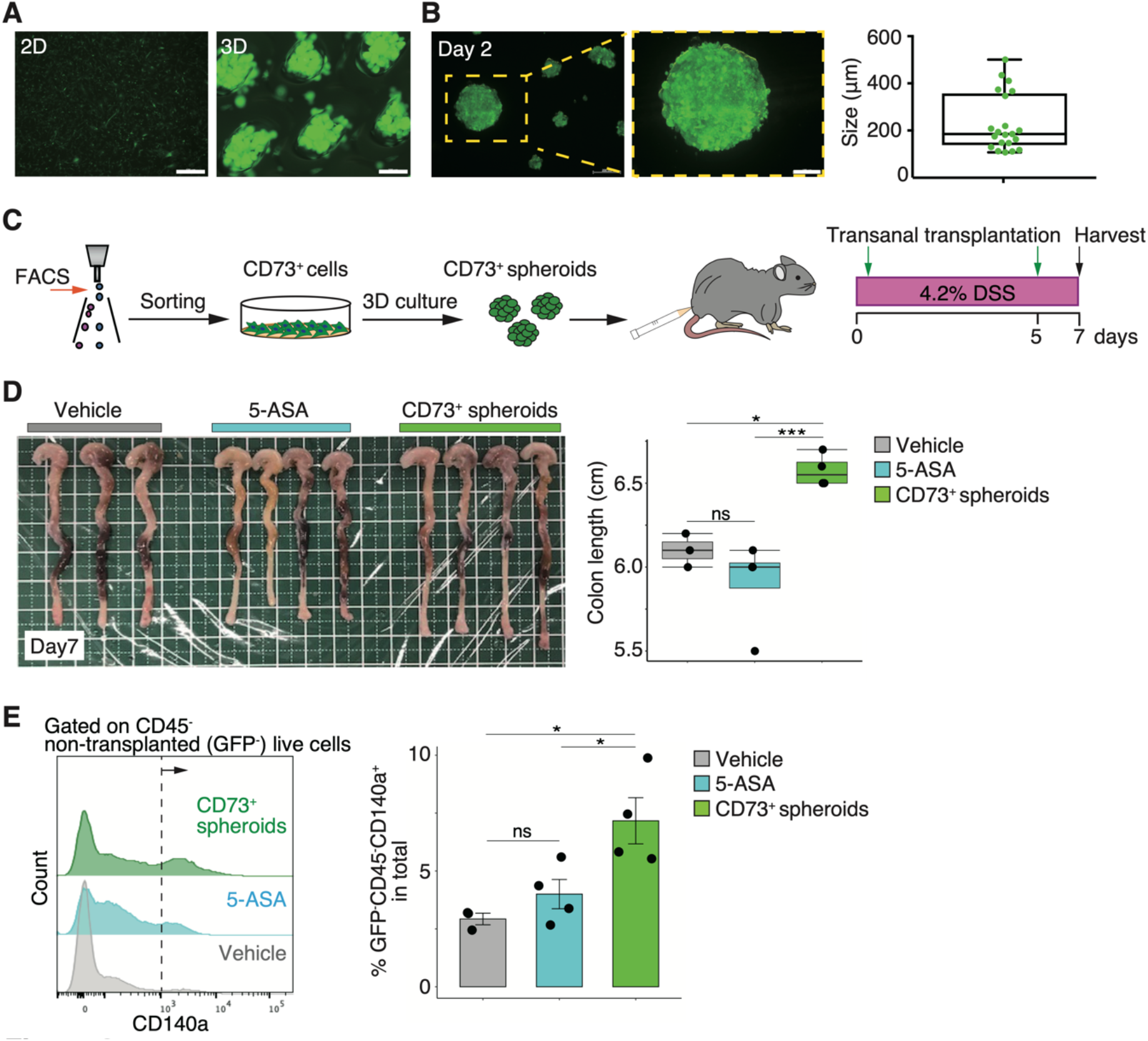
Transplantation of CD73^+^ cell spheroids onto colonic lumen prevents mucosal atrophy. A Representative images of CD73^+^ cells and 3D culture-derived spheroids. Scale bars: 2D 500 μm, 3D 200 μm. B Representative image of a single spheroid. Graph showing the spheroid size after two days of culture. Scale bar: 100 μm. C Schematic representation of the spheroid transplantation strategy via transanal transplantation. D Representative images of the colon on day 7 after DSS treatment. Graphs showing the colon length. Vehicle, PBS-treated group; 5-ASA, 5-ASA-treated group; CD73^+^ spheroids, CD73^+^ cell spheroid-transplanted group. E Histogram showing GFP^−^CD45^−^CD140a^+^ single live cells in each group indicated in the panel by FACS analysis. The dashed line denotes the threshold of the positive rate, and the graphs on the right show the quantitative data on day 7. Statistical significances were determined using Tukey’s test. Data shown as mean ± SEM. *p < 0.05; ***p < 0.005; ns, not significant.

On day 7 after DSS administration, CD45^+^CD11b^+^F4/80^+^ macrophages increased in colon tissues (**Appendix Figs S6A and B**), and we further investigated the effect of CD73^+^ spheroid transplantation on immune cells. Exactly 7 days after DSS administration, the population of these macrophages showed a declining trend in the CD73^+^ spheroid-transplanted group (mean: PBS 6.3%, 5-ASA 5.5%, and CD73^+^ spheroids 4.3%). The proportion of CD45^+^CD11b^int^F4/80^+^ macrophages also showed a declining trend in both the 5-ASA-treated and CD73^+^ spheroid-transplanted groups (mean: PBS 7.4%, 5-ASA 2.8%, and CD73^+^ spheroids 3.1%; **Appendix Figs S7A and C**). In contrast, the population of CD45^+^CD11b^−^ lymphocytes tended to increase in the CD73^+^ spheroid-transplanted group (mean: PBS 2.6%, 5-ASA 2.5%, and CD73^+^ spheroids 7.9%; **Appendix Figs S7B and C**).

### Three-dimensional spheroids enhance the properties of CD73^+^ cells

To determine the mechanism underlying the reduction of colitis in the CD73^+^ cell spheroid-transplanted group, we characterized different cell forms in terms of gene and protein expression (**Fig 4A**). We first examined protein expression using FACS analysis and found that CD73 expression was significantly higher in 3D cultures (p = 0.049; **Fig 4B**). Further analysis of other MSC markers revealed no difference in the expression of CD44 and CD90, but the expression of CD140a and Sca-1 was significantly lower in 3D cultures than in 2D cultures (p = 0.02 and p = 0.01, respectively; **Appendix Figs S8A and B**).

**Figure 4.**
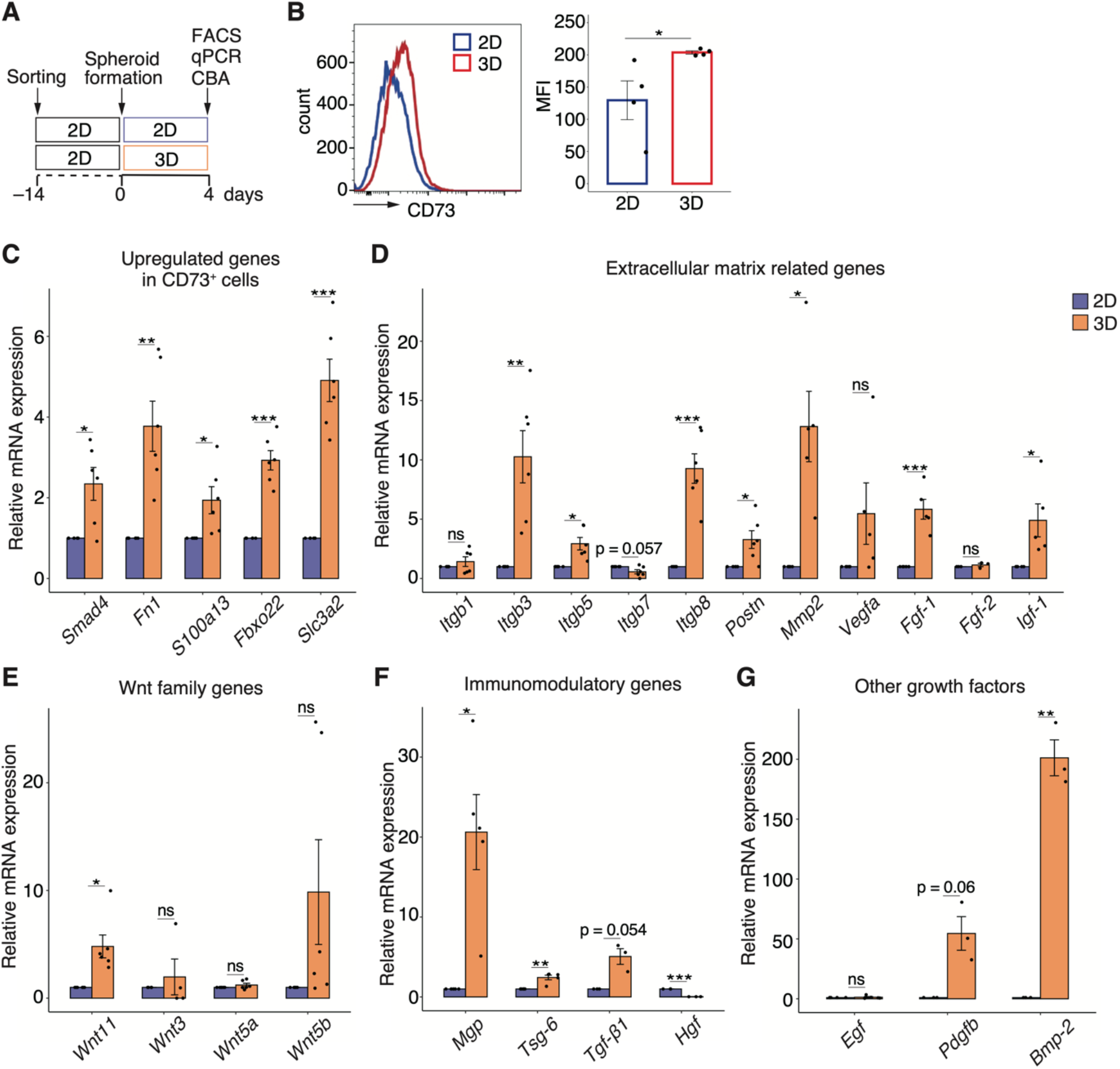
Three-dimensional spheroids enhance properties of CD73^+^ cells. A Schematic representation of the comparative analysis indicated in the panel for 2D and 3D cultures of CD73^+^ cells. B Histogram showing the frequency of CD73^+^ cells after four days of culture as determined using FACS analysis. The graph shows the quantification of mean fluorescence intensity (MFI). C–G Relative mRNA expression of the upregulated genes identified by transcriptome analysis in CD73^+^ cells (C), extracellular matrix-related genes (D), Wnt family genes (E), immunomodulatory genes (F), and interest growth factors (G). Statistical significances were determined using Welch’s t-test. Data shown as mean ± SEM. *p < 0.05; **p < 0.01; ***p < 0.005; ns, not significant.

Next, we analyzed the genes that were significantly upregulated in CD73^+^ cell populations compared with those in cMSCs (**Fig 2A**). The relative expressions of *Smad4, Fn1, S100a13, Fbxo22*, and *Slc3a1* were significantly higher in CD73^+^ spheroids than in 2D cells (**Fig 4C**). Moreover, because gene sets related to ECM, such as integrin signaling and angiogenesis, were enriched in CD73^+^ cells (**Fig 2B**), we investigated the expression of genes involved in the same and found that *Itgb3, Itgb5, Itgb8*, periostin (*Postn*), *Mmp2, Fgf-1*, and *Igf-1* were significantly upregulated in 3D cultures (**Fig 4D**). Although Wnt family genes were slightly enriched in CD73^+^ cells, *Wnt11* expression was significantly upregulated in spheroids (**Fig 4E**). We investigated gene expression by focusing on the immunomodulatory mechanisms of MSCs. mRNA expression of matrix Gla protein (*MGP*) and tumor necrosis factor (TNF) α-stimulated gene 6 (*TSG-6*) was significantly upregulated, whereas hepatocyte growth factor (*HGF*) expression was significantly downregulated in 3D cultures (**Fig 4F**). The expression of another growth factor, *Bmp-2*, which promotes osteogenic differentiation, was significantly upregulated in 3D cultures (**Fig 4G**).

To further characterize the secretory profile of CD73^+^ cell spheroids, we carried out a flow cytometric bead array (CBA) of various cytokines related to inflammatory signatures that were downregulated in CD73^+^ cells compared with cMSCs (**Fig 2A**). Comprehensive analysis of culture supernatants revealed that the levels of CCL2, CCL5, CXCL1, G-CSF, GM-CSF, and interleukin (IL)-6 were significantly lower in 3D cultures than in 2D cultures (**Fig 5**). In contrast, IL-9 and IL-17A levels tended to increase in 3D cultures (**Fig 5**).

**Figure 5.**
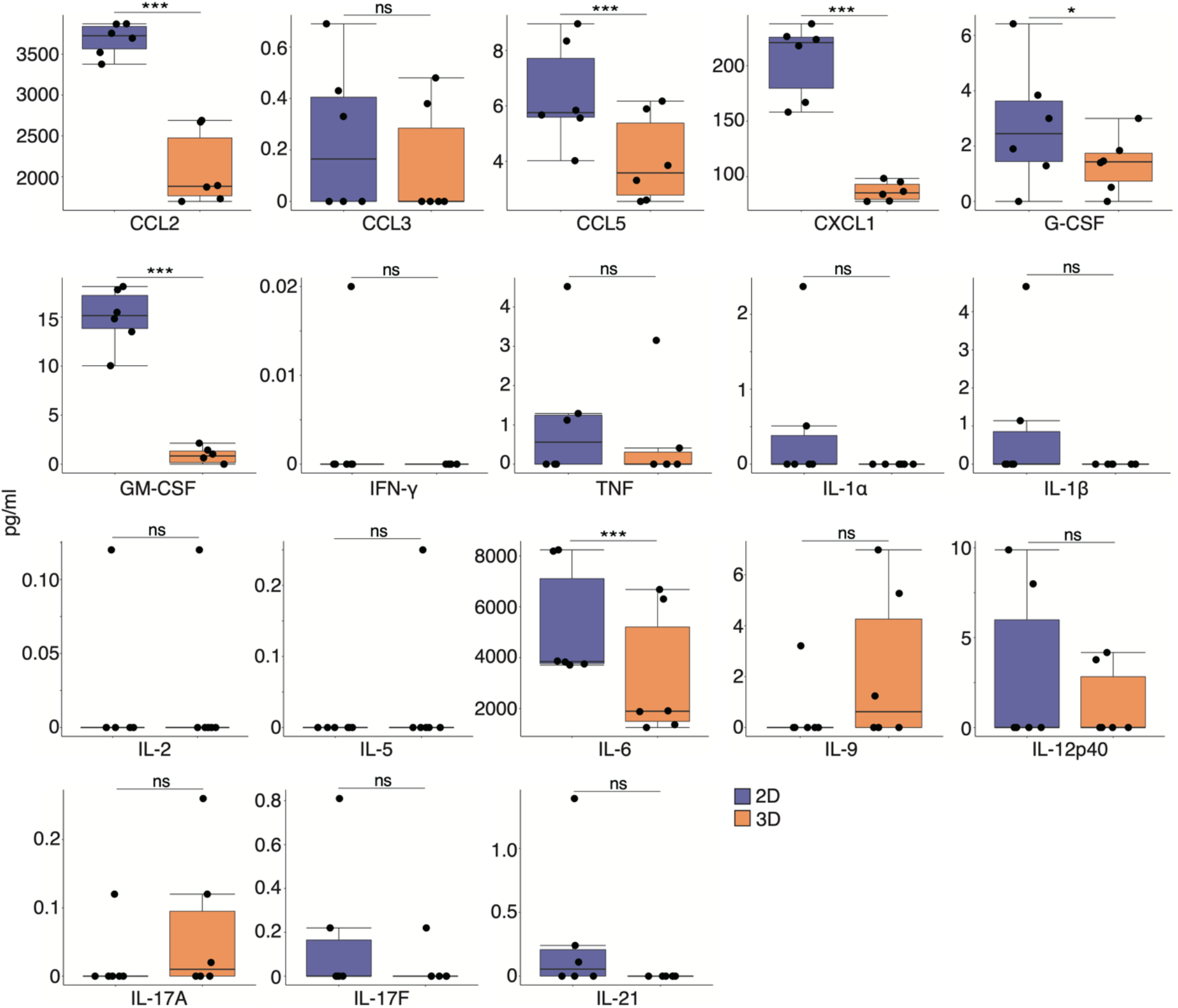
Inflammatory cytokine levels are downregulated by secretory factors of CD73^+^ cell spheroids. Cytokines in the culture supernatants of 2D and 3D cells were quantified using a cytometric bead array. Statistical significance was determined using Welch’s t-test. Data shown as mean ± SEM. *p < 0.05; ***p < 0.005; ns, not significant.

### Secretory factors of CD73^+^ cell spheroids induce alteration of ECM remodeling in fibroblasts

Although we showed that transanal transplantation of CD73^+^ spheroids attenuated DSS-induced colonic atrophy and increased tissue-resident fibroblasts (**Figs 3D and E**), the molecular mechanism remains unclear. Motivated by the fact that stromal cells, including fibroblasts with myofiber features, are associated with UC (Pasztoi & Ohnmacht, 2022), we investigated the effects of alterations in fibroblasts by secretory factors of CD73^+^ cell spheroids. To this end, we prepared conditioned media (CM), without fetal bovine serum (FBS), of CD73^+^ 2D cells and spheroids; cultured fibroblasts in regular medium supplemented with 50% CM; and analyzed cell proliferation, contraction, and gene expression at each time point (**Fig 6A**). On day 14, growth assay showed significantly lower proliferation of fibroblasts cultured in CM of 3D cultures than of those cultured in CM of 2D cultures (fold change 0.78, p < 0.01; **Fig 6B**). Furthermore, the rate of cell death remained unchanged (**Fig 6C**), suggesting that the secretory factors of the culture supernatant affected the proliferative capacity. In contrast, cell contraction assay did not show any difference between CM of 2D and 3D cultures (**Fig 6D**).

**Figure 6.**
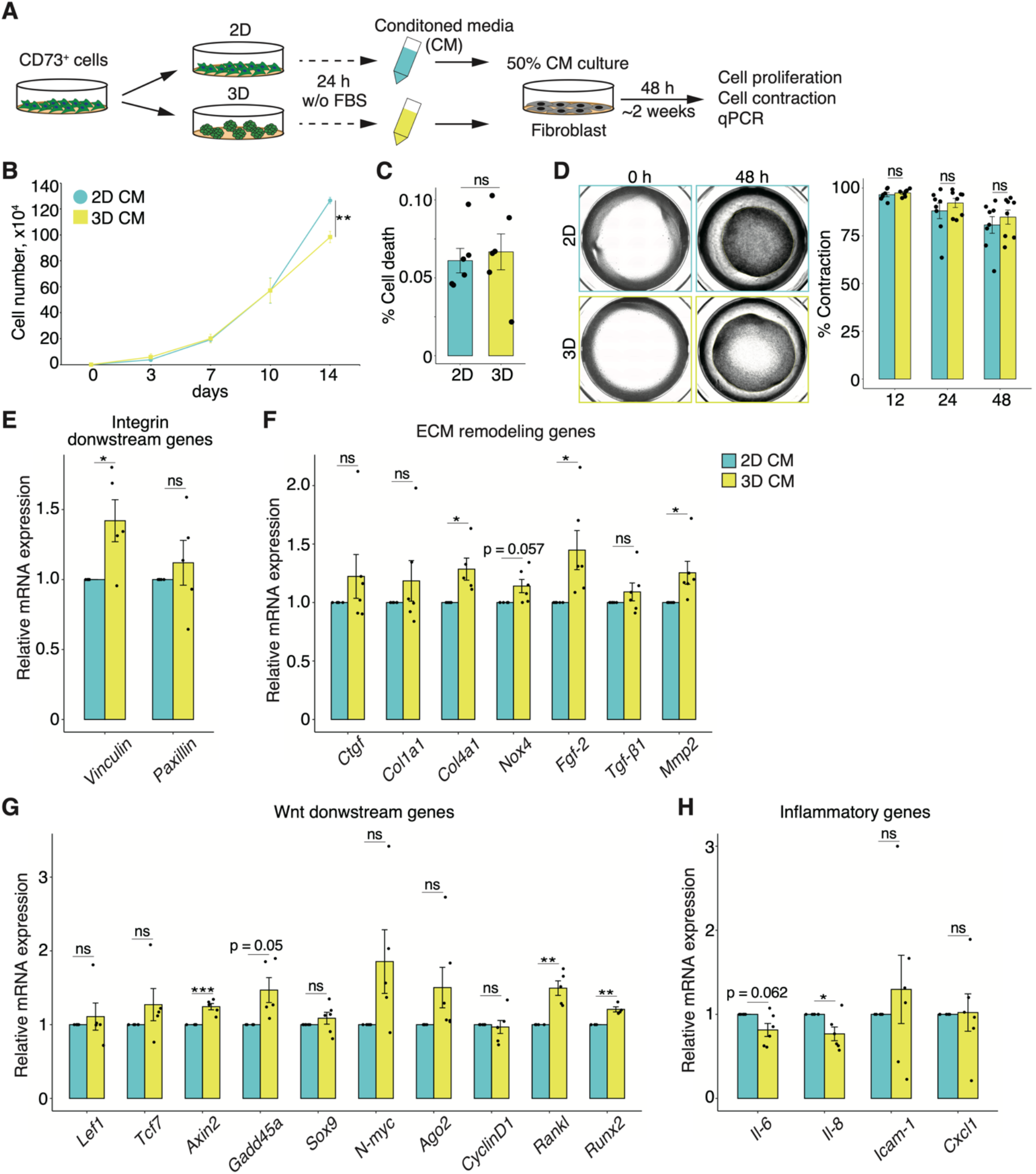
Secretory factors of CD73^+^ cell spheroids induce alteration of ECM remodeling in fibroblasts. A Schematic representation of each conditioned medium (CM) assay on fibroblasts. B Graph showing the number of cells at each time point. C Percentage of cell death at 14 days of culture. D Representative images of cell contraction assay after 48 hours. The graph shows the contraction rate at each time point. E–H Relative mRNA expression of integrin downstream genes (E), ECM remodeling genes (F), Wnt downstream genes (G), and inflammatory genes (H) in fibroblasts after 48 hours of each CM culture. Statistical significances were determined using Welch’s t-test. Data shown as mean ± SEM. *p < 0.05; **p < 0.01; ***p < 0.005; ns, not significant.

Finally, we investigated the effect of culture in CM of 3D and 2D cultures on fibroblasts. The expression of *VCL*, which encodes vinculin and is an integrin-downstream gene, and ECM remodeling-related genes *Col4a1, Mmp2*, and *Fgf-2*, was upregulated in fibroblasts cultured in CM of CD73^+^ cell spheroids (**Figs 6E and F**). Moreover, the expression of Wnt-downstream genes, such as *Axin2, Gadd45a, Rankl*, and *Runx2*, was upregulated in fibroblasts cultured in CM of 3D cultures (**Fig 6G**). In contrast, *Il-6* and *Il-8* expression was downregulated in fibroblasts cultured in CM of 3D cultures (**Fig 6H**). Altogether, these data suggest that the secretory factors of CD73^+^ cell spheroids change the gene expression profile of fibroblasts and specifically induce ECM remodeling and suppression of inflammation.

## Discussion

In this study, we demonstrated that CD73^+^ cells prospectively isolated from adipose tissue exhibited reduced inflammatory signature expression and elevated ECM remodeling related gene expression, including those involved in the Wnt signaling pathway, compared with conventionally heterogeneous MSCs. Furthermore, 3D culture-derived spheroids of CD73^+^ cells showed enhanced cellular characteristics and upregulated expression of immunomodulatory genes, such as *Mgp, Tsg-6*, and *Bmp-2*, compared with 2D cells (**Fig 7A**). Transanal transplantation of CD73^+^ cell spheroids increased their engraftment rate onto the colonic lumen and the proportion of tissue-resident CD140a^+^ fibroblasts and attenuated colonic atrophy in a mouse model of DSS-induced colitis. Moreover, we found that secretory factors accelerated ECM remodeling and suppressed excessive inflammation in fibroblasts (**Fig 7B**). Collectively, these data suggest the mitigation of colonic atrophy via remodeling of tissue-resident fibroblasts by the synergistic effects of secreted factors derived from engrafted spheroids of CD73^+^ cells.

**Figure 7.**
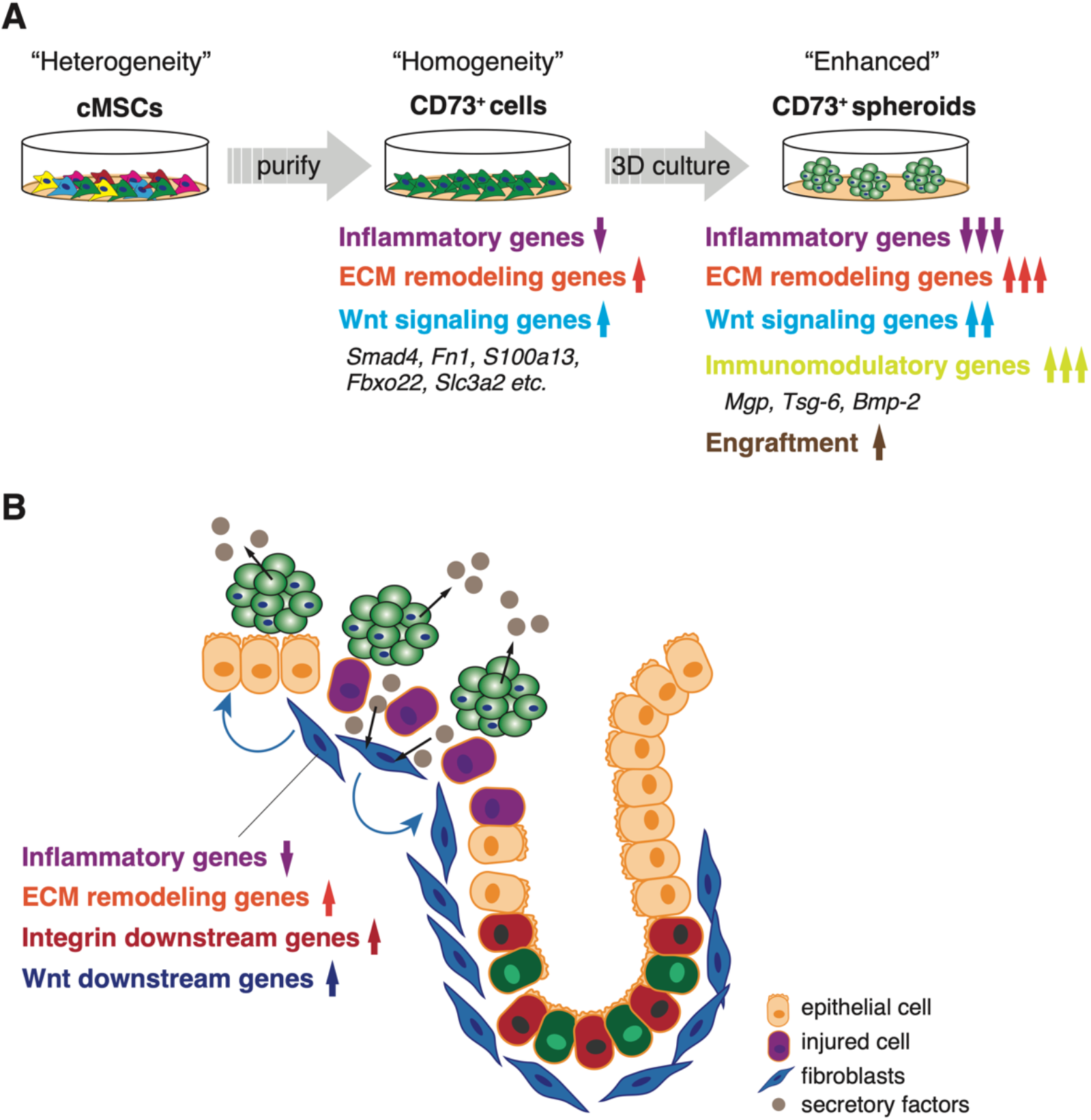
Proposed model for interaction between intestinal fibroblasts and transplanted CD73^+^ cell spheroids in DSS-induced colitis. A Compared with conventional heterogeneous MSCs, the homogeneous CD73^+^ cell population purified by prospective isolation exhibits characteristic gene expression, including downregulation of inflammatory factors and upregulation of ECM remodeling- and Wnt signaling-related genes. Moreover, 3D culture-derived spheroids of CD73^+^ cells enhance these characteristics and induce the upregulation of immunomodulatory factor expression, suggesting enhanced engraftment into colon tissues after DSS treatment. B Transplanted CD73^+^ cell spheroids attenuate colitis by modifying the phenotype of the tissue-resident fibroblasts, including downregulation of expression of inflammatory factors and upregulation of expression of ECM remodeling- and Wnt signaling-related genes.

Fibronectin, a glycoprotein characteristically expressed in CD73^+^ cells and spheroids (**Figs 2A and 4C**), is a key regulator of ECM remodeling through its involvement in cell proliferation and adhesion signaling (Bonnans *et al*, 2014). Fibronectin protein is abundant in the colonic epithelium in a steady state, but its expression is elevated in the colonic tissues of patients with IBD or DSS-induced colitis mice (Lawrance *et al*, 2017). Epithelium-derived fibronectin enhances cell adhesion and wound healing (Kolachala *et al*, 2007). SLC3A2, also known as a heavy chain of CD98 (CD98hc), regulates integrin signaling, which is pivotal for ECM remodeling (Feral *et al*, 2005; Feral *et al*, 2007). This factor contributes to integrin-dependent cell migration and protection from cell death in fibroblasts and modulates intestinal homeostasis, especially through its expression in epithelial cells under inflammatory conditions (Nguyen *et al*, 2011). Altogether, these previous studies and our findings suggest that the effectiveness of CD73^+^ cell spheroids in attenuating colonic atrophy is attributable to interaction or synergistic effect among exogenously transplanted cells, fibroblasts, and epithelial cells.

S100A13 and FBXO22 are involved in angiogenesis (Ambartsumian *et al*, 2001; Zheng *et al*, 2020), which is also a crucial process in wound healing that regulates ECM remodeling (Stupack & Cheresh, 2002). In particular, S100A13, a small Ca^2+^-binding protein, is thought to regulate co-secretion of FGF-1, a potent angiogenic factor, or they are though to enhance each other’s expression (Hayrabedyan *et al*, 2005; Landriscina *et al*, 2001). *S100a13* and *Fgf-1* expression was upregulated in CD73^+^ cell spheroids (**Figs 4C and D**), suggesting that S100A13 and FGF-1 may work in tandem to promote angiogenesis at injured sites.

Given the paracrine effects of engrafted CD73^+^ cell spheroids, we focused on secretory factors, including Wnt signaling molecules. Previous studies have shown that fibroblasts that release Wnts and POSTN are essential for supporting intestinal stem cell renewal (Degirmenci *et al*, 2018). However, upon inflammation, such as in colitis, activated fibroblasts that exhibit inflammatory signatures impair epithelial regeneration and further exacerbate pathogenesis (Kinchen *et al*, 2018; Smillie *et al*, 2019). We speculate that CD73^+^ cell spheroids with elevated expression of Wnt signaling pathway-related genes and *POSTN* replace fibroblasts that maintain intestinal homeostasis at the inflamed site, suppress inflammation, and promote early repair or protect the niche, including fibroblasts present in the steady state. Furthermore, transplantation of CD73^+^ cell spheroids increased the number of CD140a^+^ fibroblasts present in the colonic tissue (**Fig 3E**), and secretory factors released by CD73^+^ cell spheroids led to activation of the Wnt signaling pathway and downregulation of inflammatory factors in fibroblasts (**Figs 6G and H**). However, the effect of CD73^+^ cell spheroids on inflammation-associated fibroblasts remains unclear. Recent findings indicate that IL-11^+^ fibroblasts promote colorectal cancer and acute colitis (Nishina *et al*, 2021). Using *Il11* reporter mouse to perform CD73^+^ cell transplantation experiments will clarify its effects on inflammatory fibroblasts.

During the wound healing process, an inflammatory state has both negative and positive impacts (Martin & Leibovich, 2005). Infiltration of immune cells, such as macrophages and neutrophils, into the injured site promotes tissue repair (Sorokin, 2010). In the present study, we used 5-ASA, which is prescribed to more than 80% of patients with UC as a first-line treatment (Chapman *et al*, 2020), as a positive control to ascertain the therapeutic efficacy of CD73^+^ cell spheroids. The benefits of 5-ASA are mainly due to its anti-inflammatory effects (Abdu-Allah *et al*, 2016). However, CD73^+^ cell spheroids did not appear to improve colon length or affect fibroblasts during the acute phase of DSS-induced colitis (**Figs 3D and E**). Moreover, the effects of 5-ASA treatment on lymphocytes and macrophages differ from those of CD73^+^ spheroid transplantation (**Appendix Fig S7**), suggesting that the mode of action is different. MSCs are known to act on a myriad of immune cells, and this immunomodulatory property is a pivotal function in clinical applications (Lee & Kang, 2020; Wu *et al*, 2020). TSG-6, an immunomodulatory factor, improves myocardial infarction through its anti-inflammatory effects (Lee *et al*, 2009). Furthermore, MSC aggregation augments TSG-6 expression (Bartosh *et al*, 2010), which is consistent with our results (**Fig 4F**). In addition, MGP, whose expression was upregulated in CD73^+^ cell spheroids (**Fig 4F**), reportedly ameliorates pathogenesis in a mouse model of Crohn’s disease through its MSC-derived immunomodulatory function that suppresses T cell proliferation and cytokine production (Feng *et al*, 2018). Overall, these results suggest that anti-inflammatory drugs, such as 5-ASA, may compromise tissue repair due to their excessive inhibitory effects, whereas CD73^+^ cell spheroids, which express high levels of immunomodulatory factors, may regulate the immune balance, leading to earlier recovery or diminished colonic atrophy.

The present study has several limitations. First, it is unknown whether CD73^+^ cell spheroid transplantation has potential side effects. Inhibition of CD73 activity or gene silencing in tumor cells is reported to attenuate tumorigenesis (Jin *et al*, 2010; Stagg *et al*, 2010). Additionally, CD73 is highly expressed in solid tumors and is reported to facilitate colitis-associated tumorigenesis (Liu *et al*, 2020; Roh *et al*, 2020). These findings suggest that transplantation of CD73^+^ cells could lead to tumor formation. However, the promotion of colitis-associated tumorigenesis by CD73 reported by Liu *et al* was observed in a chronic inflammatory model generated following three cycles of treatment with azoxymethane, a genotoxic colonic carcinogen, and DSS administration for 3 weeks (Liu *et al*, 2020), which is different from the acute colitis model used in the present study. Moreover, transplanted MSCs disappear after temporary engraftment (Kean *et al*, 2013), but we will further investigate whether CD73^+^ cell spheroids remain at the injured site for a long time using longitudinal analysis, considering the potential clinical application of MSC transplantation. Second, the function of CD73 remains unclear. CD73 is a glycosyl phosphatidylinositol anchored membrane protein and one of the ectonucleotidases that metabolize extracellular adenosine triphosphate (ATP) (Kepp *et al*, 2021). CD73 converts 5′-adenosine monophosphate (AMP) to adenosine, which contributes to immunosurveillance in various tumor microenvironments (Kepp *et al*, 2021). Our data also showed that expression of gene sets related to metabolism, such as “GTP biosynthetic process” and “UTP biosynthetic process,” is enriched in CD73^+^ cells (**Fig 2A**), suggesting that the transplanted CD73^+^ cell spheroids have a distinct adenosine-mediated role independent of the ECM remodeling pathway reported in the present study. Adenosine in colitis regulates inflammatory responses, such as epithelial hyperpermeability, but it elicits different phenotypes depending on the receptor subset (Bahreyni *et al*, 2018). In addition, CD73 and adenosine receptors are widely expressed on tumor cells and immune cells, including regulatory T cells, which play a role in suppressing inflammation, and play context-dependent roles in various cell types (Roh *et al*, 2020). Specifically, primary colorectal cancers associated with IBD are adenocarcinomas consisting of epithelial cell-derived tumor cells (Tanaka *et al*, 2003), suggesting that CD73 expressed in MSCs has a distinct function. Clarifying the function of CD73 in MSCs is essential for clinical application of CD73^+^ cells.

In summary, we demonstrated an appropriate transplantation method based on the characteristics of a uniform CD73^+^ cell population and revealed the interaction between transplanted cells and tissue-resident fibroblasts as one of the mechanisms of colitis repression (**Figs 7A and B**). Our work opens new avenues for treating UC and other inflammation-based diseases. Our findings suggest a potential role for CD73^+^ cell spheroids as a source of transplanted cells or secreted factors in the clinical application of MSCs.

## Materials and Methods

### Experimental animals

Male C57BL/6 NJcl (wild-type) mice (10–12 weeks old, average weight 25.16 g) were obtained from CLEA Japan, Inc. (Tokyo, Japan), and male C57BL/6-Tg (CAG-EGFP) mice (6–8 weeks old, average weight 19.78 g) were obtained from Japan SLC, Inc. (Hamamatsu, Japan). Donor cells used for transplantation were derived from CAG-EGFP mice to distinguish them from the wild-type recipient mice. All experimental animal procedures were approved by the Animal Committee of Tokyo Medical and Dental University (#A2019-307A) and Juntendo University School of Medicine (#1477). All methods were conducted in strict accordance with the approved guidelines of the Institutional Animal Care Committee. All mice were housed in cages in a specific pathogen-free room under 12-h light/dark cycles and received water and food ad libitum.

### DSS-induced colitis model

For the induction of colitis, 3.0% or 4.2% w/v DSS solution (MP Biomedicals, Santa Ana, CA, USA) was supplemented in drinking water and administered to wild-type mice for 5 or 7 days, with a change of water every three days. After DSS treatment, the mice were given regular water. Changes in body weight were measured. Mice were anesthetized with isoflurane and sacrificed for tissue sampling. In the present study, two different DSS concentrations (3.0% and 4.2%) were used to account for variations by lot. The concentration at which weight loss was observed in more than 90% of the individuals through pilot experiments was adopted for each lot of DSS.

### Isolation and intravenous administration of CD73-positive cells

CD73^+^ cells were isolated from the subcutaneous fat tissue of mice, as previously described (Suto *et al*, 2017). Cells were washed with Hank’s balanced salt solution (HBSS, Nacalai Tesque, Kyoto, Japan) and 4% FBS (Thermo Fisher Scientific, Waltham, MA, USA) before staining with a fluorescence-conjugated antibody cocktail for 30 min on ice. The following antibodies were used: Ter119 (TER119), CD45 (30-F11), CD31 (390), and CD73 (TY11.8) (BD Biosciences, Franklin Lakes, NJ, USA). CD73^+^ cells were defined as Ter119^−^CD45^−^ CD31^−^CD73^+^ live cells and sorted on a FACS AriaII (BD Biosciences). All experiments were analyzed using the FlowJo software (TreeStar, Ashland, OR, USA). CD73^+^ cells were cultured in an MSC medium, as previously described, and were never passaged more than five times (Suto *et al*, 2017). After expanded culture, CD73^+^ cells (3.0 × 10^5^ cells in 300 μL of PBS) were administered to the mice through intravenous injection 1 and 5 days after DSS treatment.

### Spheroid formation and transanal transplantation

Exactly 14 days after sorting, CD73^+^ cells were cultured for two days in EZSPHERE^®^ (AGC Techno Glass, Shizuoka, Japan) to generate spheroids. Spheroids were collected and suspended in PBS. Cells (1.0 × 10^6^ cells in 200 μL of PBS) were transplanted via enteral administration 0 and 5 days after DSS treatment. Transanal transplantation was performed as previously described with slight modifications (Fukuda *et al*, 2014). The cell suspension was infused into the murine colonic lumen using a thin, flexible catheter (diameter: 2.1 mm) or stainless steel needle (diameter: 1.9 mm). After transplantation, the mice were housed as usual, and colon tissues were collected and analyzed by flow cytometry. Control mice received an equal volume of PBS or a solution of 0.01 mg/mL 5-ASA (Zeria Pharmaceutical Co., Ltd., Tokyo, Japan).

### Flow cytometry

For analysis of colonic tissue, after measuring the colon length, the distal colon, 2 cm above the rectum, was cut longitudinally and washed with PBS. The tissue was minced and digested with 2 mg/mL collagenase (FUJI-FILM Wako Pure Chemical, Osaka, Japan) solution containing 10 μM DNase I (Sigma-Aldrich, St. Louis, MO, USA) in Dulbecco’s modified Eagle medium (DMEM)-GlutaMAX (Thermo Fisher Scientific) with shaking for 1 h at 37 °C. The digested tissue was filtered through a 70-μm cell strainer (CORNING, Corning, NY, USA), pelleted by centrifugation at 1,800 g for 5 min at 4 °C, and resuspended in HBSS containing 2% FBS, 10 nM HEPES, and 1% penicillin/streptomycin (Thermo Fisher Scientific). The cell suspensions were stained with an antibody cocktail for 30 min on ice. The following antibodies were used: CD45 (30-F11) from BD Biosciences and CD11b (M1/70) and F4/80 (BM8) from BioLegend (San Diego, CA, USA). The following antibodies were used for analysis of MSC markers: CD73 (TY11.8), CD44 (1M7), Sca-1 (E13-161.7), and CD140a (APA5) from BD Biosciences, and CD90.2 (30-H12) from BioLegend. Data acquisition was performed using FACS AriaII (BD Biosciences), and all analyses were carried out using FlowJo software (TreeStar).

### Collection of CM

CD73^+^ cells of 2D or 3D cultures were initiated with the same number (1.0 × 10^7^) of cells and cultured in an MSC medium for 2 days. The MSC medium was replaced with fresh MSC medium without FBS before sampling and collected 24 h later.

### CBA

The expression level of cytokines in the CM was determined using CBA (BD Biosciences), according to the manufacturer’s instructions. All samples were diluted 1:2 and analyzed in duplicate. All data were analyzed on a BD FACS Verse Flow Cytometer (BD Biosciences) and FCAP Array Software v3.0 (BD Biosciences).

### Cell proliferation and contraction assays

For growth curves, murine embryonic fibroblasts (1.0 × 10^3^ cells) were seeded in 48-well plates and cultured in DMED medium (Nacalai Tesque) with 10% FBS, 1% penicillin/streptomycin, and 2 mM L-glutamine (Nacalai Tesque) supplemented with 50% CM after 1 day for up to 2 weeks, and the number of live and dead cells was measured at the indicated times. A collagen-based cell contraction assay of murine embryonic fibroblasts (1.0 × 10^5^ cells) was performed in 48-well plates, according to the manufacturer’s instructions (Cell Biolabs, Inc., San Diego, CA, USA). Before releasing the stressed matrix, the culture medium was replaced with fresh medium containing 50% CM. The collagen gel size was evaluated using the BX-H4M software of the BZ-X710 microscope (Keyence, Osaka, Japan).

### Quantitative real-time polymerase chain reaction (qPCR)

Total RNA extraction and qPCR were performed as described previously (Suto *et al*, 2017). Gene expression was normalized to *Gapdh* expression. Primer sequences used for qPCR are listed in Appendix Table S1.

### Acquisition of clinical samples

Human adipose tissue samples were provided from surplus tissues from breast reconstructive and liposuction surgeries from patients without underlying diseases such as diabetes (three individuals). Human adipose-derived CD73^+^ cells and cMSCs were established from the same samples. The study protocol was approved by the Institutional Review Board (IRB) of Juntendo University Hospital (IRB #16–102). All patients provided written informed consent for the use of tissue samples in this study.

### mRNA sequencing (mRNA-seq)

CD73^+^ cells and cMSCs were isolated from human adipose tissue and cultured as previously described (Suto *et al*, 2020). Cell suspensions were incubated with APC-conjugated anti-CD73 (BioLegend) for single-color staining. To distinguish live cells from dead cells, the cells were suspended in a propidium iodide solution and sorted using FACS Aria II (BD Biosciences). All experiments were analyzed using the FlowJo software (TreeStar). Total RNA was extracted from these cells using a RNeasy Mini kit (Qiagen, Hilden, Germany) according to the manufacturer’s instructions. Extracted RNA samples were then sent to Macrogen (Tokyo, Japan). The sequencing library was prepared using the SMART-Seq v4 Ultra Low Input RNA kit (Takara, Kusatsu, Japan), and mRNA-seq on an Illumina NovaSeq 6000 (paired-end, 150 bp) was performed by Macrogen.

To generate a sequence alignment, quality control and preprocessing of FASTQ data were performed using fastp v.0.20.1 (https://github.com/OpenGene/fastp) (Chen *et al*, 2018), and the transcript was quantified using Salmon (https://combine-lab.github.io/salmon/) (Patro *et al*, 2017). Data analyses and visualizations were conducted in the RStudio environment. The GO annotation was performed using the DAVID online software (http://david.abcc.ncifcrf.gov). To compare the enriched genes between cMSCs and CD73^+^ cells, we used GSEA software (Broad Institute, Cambridge, MA, USA).

### Histological analyses

Colonic tissue was collected from mice as previously described (Fukuda *et al*, 2014). The histological score after DSS treatment was defined using the following criteria (Erben *et al*, 2014; Ernst *et al*, 2015): mucosal architecture (0–3), immune cell infiltration (0–3), muscle layer thickness (0–3), depletion of goblet cells (0–1), and crypt abscess (0–1). H&E and immunofluorescence staining were performed as previously described (Suto *et al*, 2020).

### Statistical analysis

A minimum of three independent experiments were included in each statistical analysis. Statistical significance was determined using each test indicated in the figure legends and was set at p < 0.05. Analyses were performed in the statistical programming language R version 4.0.3 (2020-10-10).

## Data availability

mRNA-seq data supporting the findings of the present study have been deposited in the Gene Expression Omnibus SuperSeries record GSE211637 (https://www.ncbi.nlm.nih.gov/geo/query/acc.cgi?acc=GSE211637).

## Acknowledgments

We thank Dr. Ryuichi Okamoto (Tokyo Medical and Dental University) for helping with the transanal transplantation procedure and Dr. Rica Tanaka (Juntendo University School of Medicine) for providing clinical samples. We also acknowledge all lab members who were involved in this project. This work was financially supported by Otsuka Holdings Co., Ltd., Japan, and the Japan Society for the Promotion of Science (JSPS)/ Ministry of Education, Culture, Sports, Science, and Technology (MEXT) KAKENHI Grant-in-Aid for Scientific Research (B) (grant number 21H03328). This work also was partially supported by the JSPS/MEXT KAKENHI Grant-in-Aid for the Promotion of Joint International Research (A) (grant number JP18KK0449), for Scientific Research (C) (grant number JP19K10024), and for Early-Career Scientists (grant number 21K15888).

## Author contributions

**DH:** Conceptualization; data curation; visualization; formal analysis; investigation; methodology; funding acquisition; writing —original draft; writing —review and editing. **NI, RO, ES, YN, AI, and LI:** Formal analysis; investigation; methodology. **YM:** Formal analysis; investigation; methodology; funding acquisition; writing —review and editing. **CA:** Conceptualization; supervision; funding acquisition; writing —review and editing.

