## Appendix for "CD73-positive cell spheroid transplantation attenuates colonic atrophy"

**Title**

### Appendix Figures

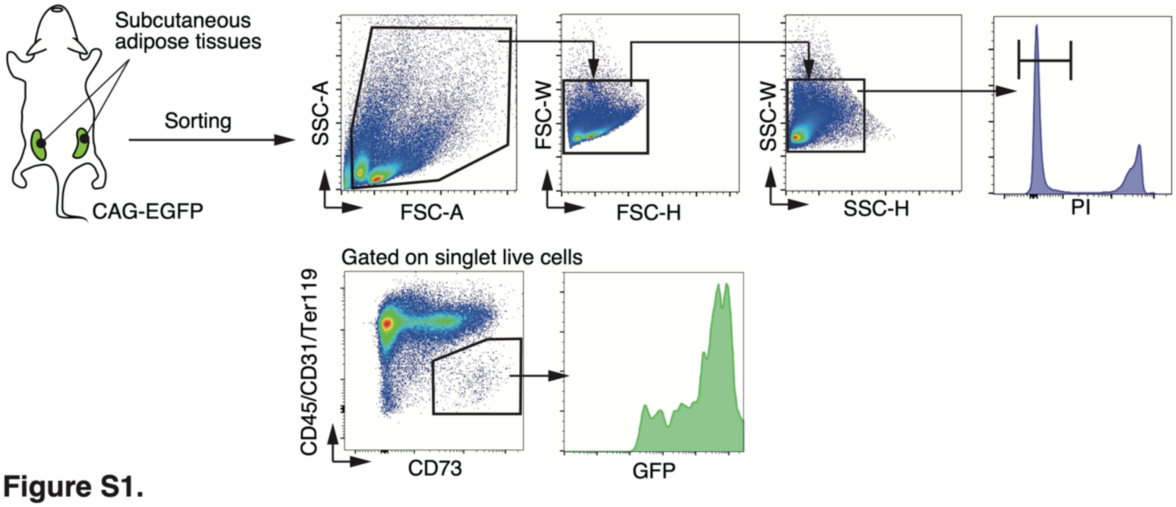


**Figure S1.**

**Appendix Figure S1 - Representative FACS plots of isolation of adipose-derived CD73$^{+}$ cells from mouse subcutaneous fat.**

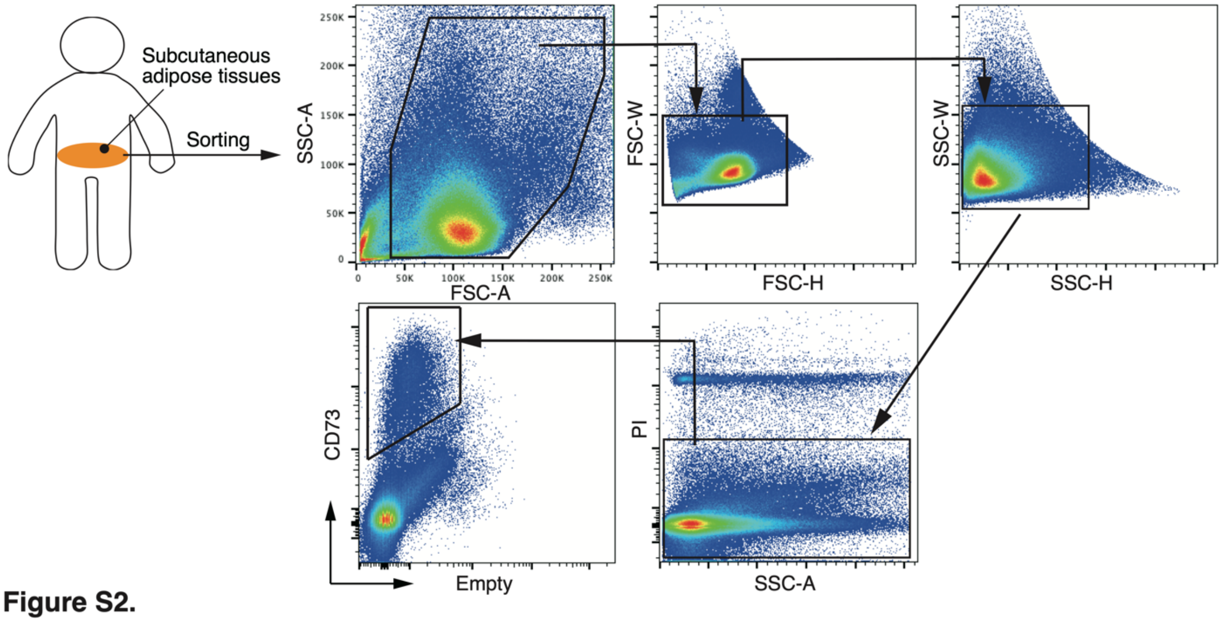


**Figure S2.**

**Appendix Figure S2 - Representative FACS plots of isolation of $CD73^{+}$ cells from human adipose tissue.**

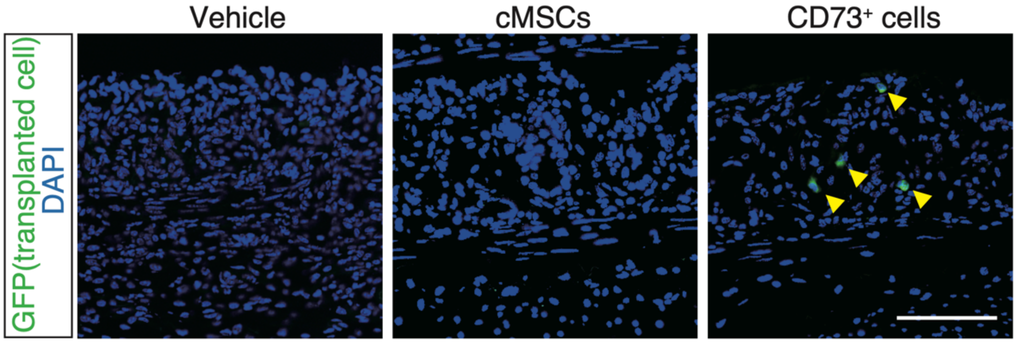


**Figure S3.**

**Appendix Figure S3 - Transplanted cell engraftment by intravenous injection.**

Representative immunohistochemistry images of $GFP^+$ transplanted cells after intravenous injection. Arrowheads indicate transplanted cells. No cell engraftment was observed in the PBS-treated and cMSC-transplanted groups. Vehicle; PBS-treated group, cMSCs; cMSC-transplanted group; $CD73^+$ cells; $CD73^+$ cell-transplanted group. Scale bar: 100 μm.

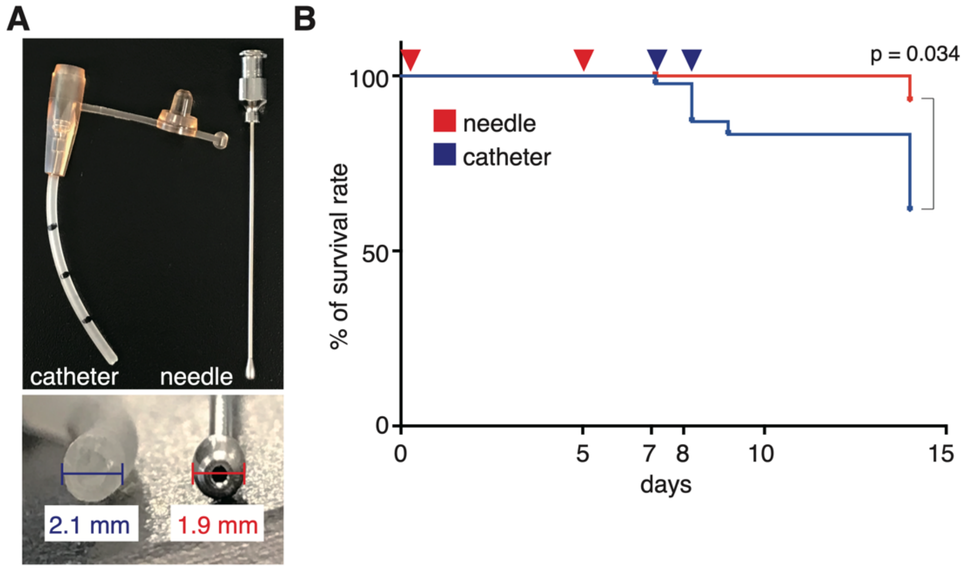


Figure S4.

**Appendix Figure S4 - Improvement of transanal transplantation method.**

A Comparison of a thin, flexible catheter and a stainless steel needle used for cell injection.

B Survival rate for each method. Arrowheads indicate transplantation day.

Statistical significances were determined using Welch's t-test.

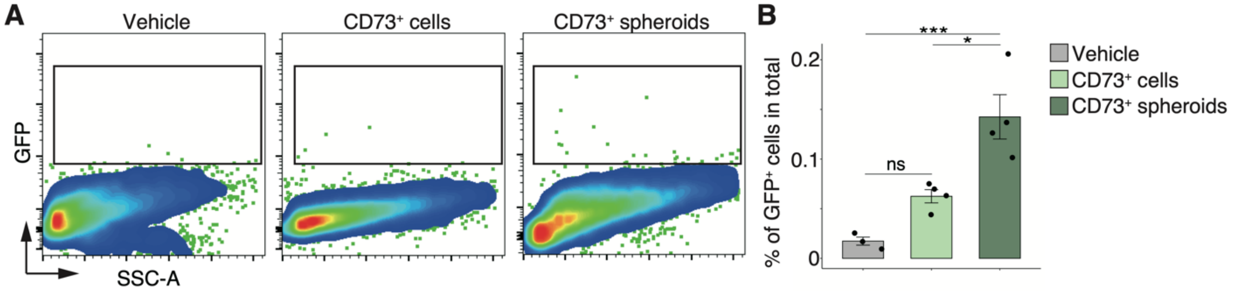


**Figure S5.**

**Appendix Figure S5 - 3D culture improves engraftment of CD73$^+$ cells into colonic tissue after DSS treatment.**

A Representative FACS plots of GFP$^+$ transplanted cells after transanal transplantation using a stainless steel needle.

B Graph showing the percentage of GFP$^+$ transplanted cells.

Vehicle; PBS-treated group, CD73$^+$ cells; CD73$^+$ 2D cell-transplanted group; CD73$^+$ spheroids; CD73$^+$ cell spheroid-transplanted group. Statistical significances were determined using Tukey's test. Data shown as mean ± SEM. *$p < 0.05$; ***$p < 0.005$; ns, not significant.

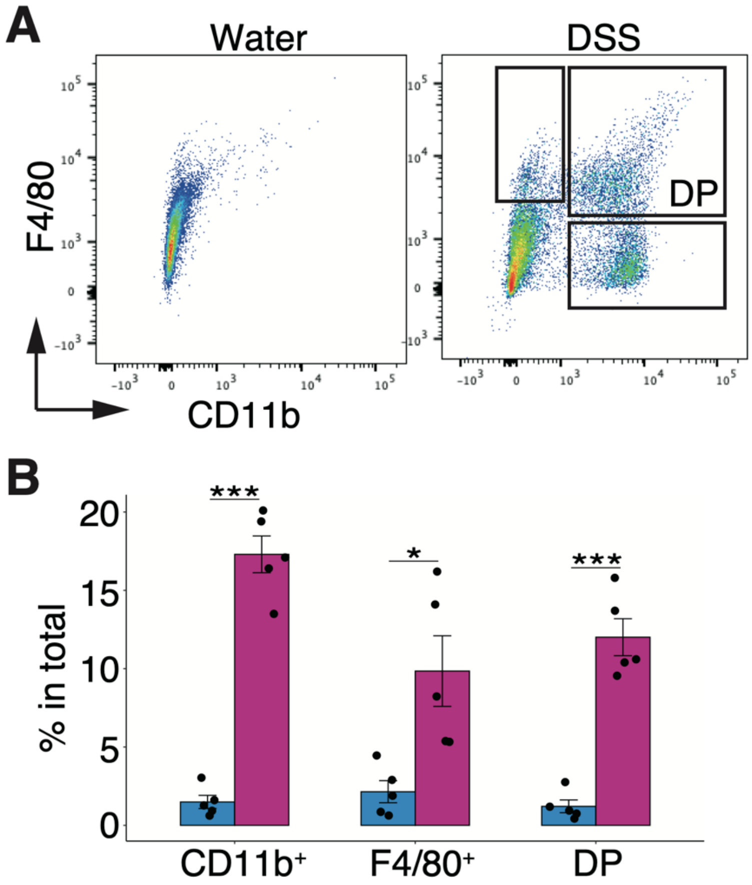


**Figure S6.**

**Appendix Figure S6 - Percentage of CD11b⁺F4/80⁺ macrophages increases in the distal colon after DSS treatment.**

A Representative FACS plots of $CD45^+$ single live cells isolated from the distal colon on day 7 after DSS treatment.

B Graph showing the percentages of each cell population indicated in the panel.

Statistical significances were determined using Welch's t-test. Data shown as mean ± SEM. *$p < 0.05$; ***$p < 0.005$. DP, double positive cell.

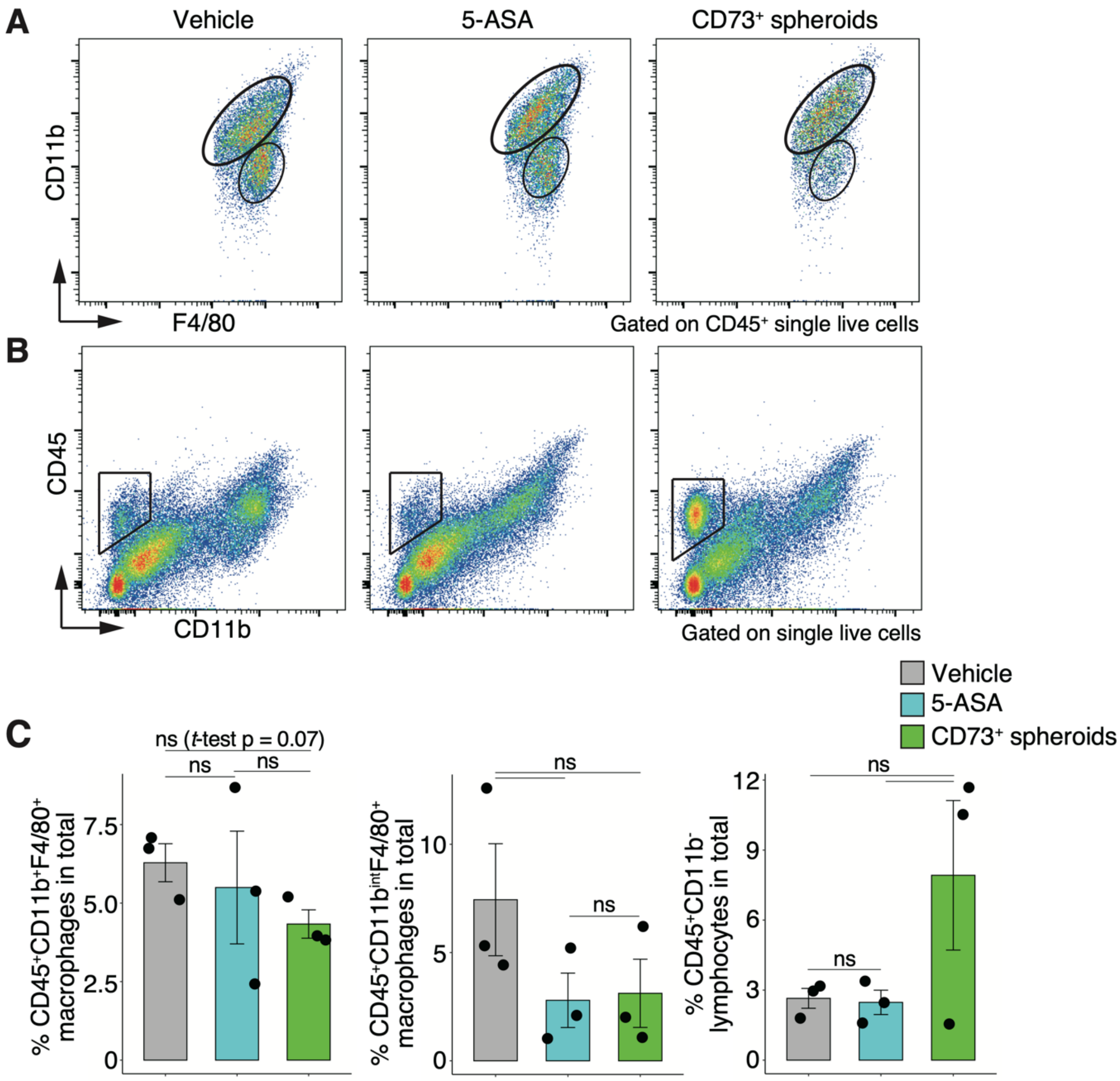


**Figure S7.**

**Appendix Figure S7 - Transanal transplantation of $CD73^+$ cell spheroid slightly affects immune cells in the distal colon.**

A, B Representative FACS plots of the colonic tissue cells gated on $CD45^+$ single live cells (A) and single live cells (B) on day 7 after DSS treatment.

C Graphs showing the percentage of $CD45^+CD11b^+F4/80^+$, $CD45^+CD11b^{int}F4/80^+$ macrophages and $CD45^+CD11b^-$ lymphocytes on day 7 after DSS treatment.

Statistical significances were determined using Tukey's test and Welch's t-test. Data shown as mean ± SEM. ns, not significant. Vehicle, PBS-treated group; 5-ASA, 5-ASA-treated group; $CD73^+$ spheroids, $CD73^+$ cell spheroid-transplanted group.

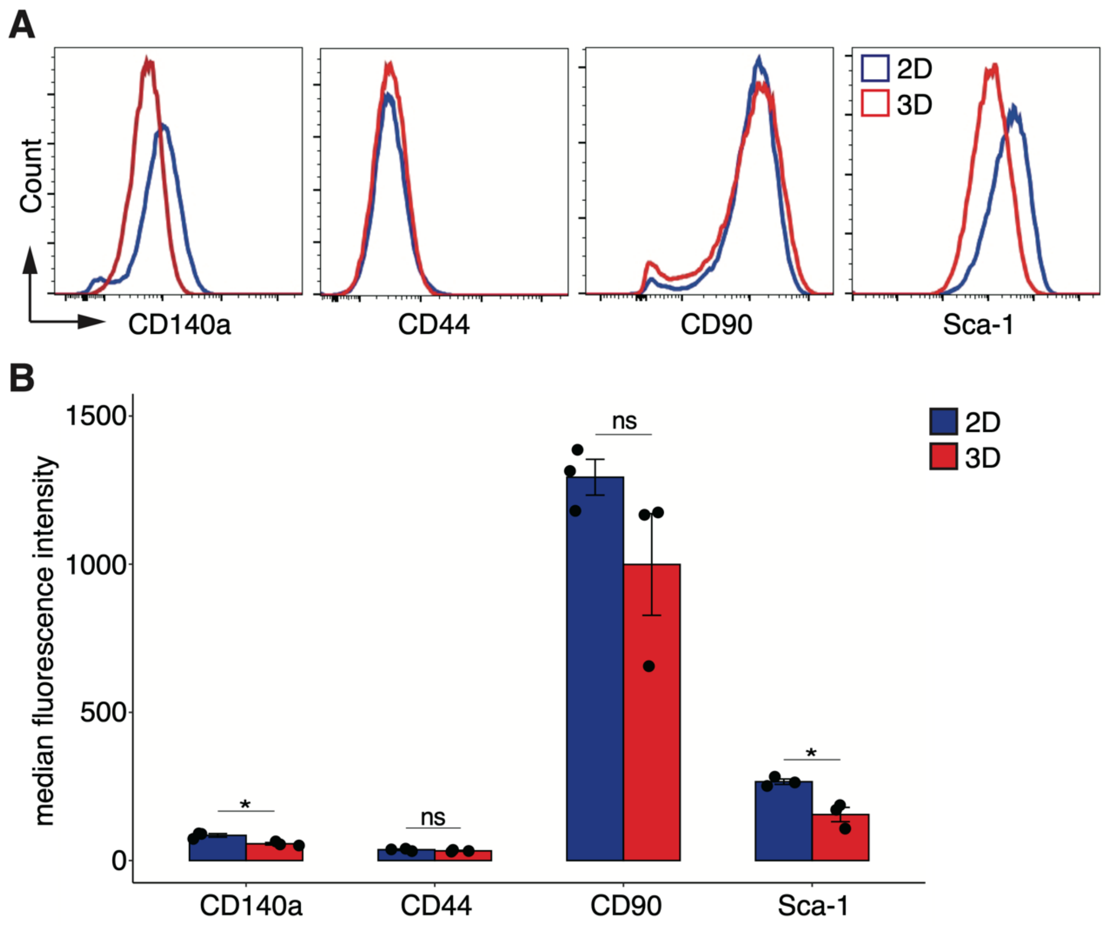


**Figure S8.**

**Appendix Figure S8 - Protein expression of MSC markers under 3D culture determined using FACS analysis.**

A Representative histograms of each MSC marker in $CD73^{+}$ 2D cells and 3D spheroids.

B Graph showing the mean fluorescent intensity of MSC markers.

Statistical significances were determined using Welch's t-test. Data shown as mean ± SEM. *$p < 0.05$. ns, not significant.

**Appendix Table S1 - Mouse qPCR primers sequences used in the present study.**

| Gene | Forward | Reverse |
|---|---|---|
| Ago2 | TCCCAGGATATGCCTTCAAA | TAAAGTGCTGGACCATGTGC |
| Axin2 | AGAAGCGACCCAGTCAATCC | TTACTCCCCATGCGGTAAGG |
| Bmp-2 | TAGATCTGTACCGCAGGCACTC | CTGCAGATGTGAGAAACTCGTC |
| Col1a1 | ATGTTCAGCTTTGTGGACCT | CAGCTGACTTCAGGGATGT |
| Col4a1 | ATTGTGGTGGCTCTGGCTGT | GCCAGGAAGTCCAGGTTCTC |
| Ctgf | CCACCCGAGTTACCAATGAC | GCTTGGCGATTTTAGGTGTC |
| Cxcl1 | CAGCCACCCGCTCGCTTCTC | AGGGAGCTTCAGGGTCAAGG |
| CyclinD1 | TGGTGAACAAGCTCAAGTGG | TGGAAAGAAAGTGCGTTGTG |
| Egf | CCCGAACTTCTCAAACAGAGG | CCATCAGGAAGCAGAACAAATC |
| Fbxo22 | GACAGTTCTCTACATGGCAGATT | ACAATTCCCGGGGTCACA |
| Fgf-1 | AAAAGCCCAAACTGCTCTACTG | GTATAAAAGCCCTTCGGTGTCC |
| Fgf-2 | AAGCGGCTCTACTGCAAGAAC | CATAGCAAGGTACCGGTTGG |
| Fn1 | GCTTTGACAAATACACTGGGAAC | CATTTCCCAGACACAGACACTC |
| Gadd45a | GCAGAGCAGAAGACCGAAAG | GCAGGCACAGTACCACGTTA |
| Gapdh | AAAGGGTCATCATCTCCGCC | CTCGTGGTTCACACCCATCA |
| Hgf | AGTGTGCCAACAGGTGTATCAG | GTCCCTTTATAGCTGCCTCCTT |
| Icam-1 | GAGAGTGGACCCAACTGGAA | GGGTGAGGTCCTTGCCTACT |
| Igf-1 | CAGCTGGACCAGAGACCCTTT | ATCACAGCTCCGGAAGCAAC |
| Il-6 | GGATACCACTCCCAACAGACC | TCCAGTTTGGTAGCATCCATC |
| Il-8 | TCAAGAGCTACGATGTCTGTGT | GGCCAACAGTAGCCTTCACC |
| Itgb1 | CCTTCAATTGCTCACCTTGTTC | CGATGATTAGCTGGATCACATTAC |
| Itgb3 | CCACACGAGGCGTGAACTC | CTTCAGGTTAC ATCGGGGTGA |
| Itgb5 | GAAGTGCCACCTCGTGTGAA | GGACCGTGGATTGCCAAAGT |
| Itgb7 | ACCTGAGCTACTCAATGAAGGA | CACCGTTTTGTCCACGAAGG |
| Itgb8 | CTGAAGAAATA CCCCGTGGA | ATGGGGAGGCATACAGTCT |
| Lef1 | GTCAGACAAGCCCCGTCCT | GCTGTTCATATTGGGCATCATT |
| Mgp | GCTCCCTCTGGCCATCCT | GCTTTAGCTCGCCACCTCT |
| Mmp2 | ACCAGAACACCATCGAGACC | TACTTTTAAGGCCCGAGCAA |
| N-myc | TCCTGGGAACTGGGTTGGAG | GACCGCCGAAGTAGAAGTCA |
| Nox4 | CTTTTCATTGGGCGTCCTC | GGGTCCACAGCAGAAAACTC |
| Paxillin | ACTACTGCAACGGACCCATC | TAGTGGACCTCACAGTACGG |
| Pdgfb | CTCCATCCGCTCCTTTGATG | GATCGATGAGGTTCCGAGAGA |
| Postn | CTCAGCACTACTCCGATGTCTC | TTAACCATGTGGCTGTGTAAGG |
| Rankl | AAACGCAGATTTGCAGGACT | ACATCCAACCATGAGCCTTC |
| Runx2 | GCGTCAAACAGCCTCTTCAG | GTTGTTGCTGTTGCTGTTGT |
| S100a13 | TGGTCTCTACTTTCTTCACCTTTG | TCACATCCAAGGTCTTCATCTTT |
| Slc3a2 | CAGCTCCTACCTGTCAAATTCC | TGGTAGAGTCGGAGAAGATGGT |
| Smad4 | AGGACATTCGATTCAAACCATC | TTTCAAAGTAAGCAATGGAGCA |
| Sox9 | AAGAACAAGCCACACGTCAA | CGTTCTTCACCGACTTCCTC |
| Tcf7 | ATCCTTGATGCTGGGATCTG | CTTCTCTGCCTTGGGTTCTG |
| Tgf-β1 | GTCAGACATTCGGGAAGCAGT | GGTCAGCAGCCGGTTACCAAG |
| Tsg-6 | CTGGCAGATACAAGCTCACCTAC | GGGTATCCGACTCTACCCTTG |
| Vegfa | CCATGAACTTTCTGCTCTCTTG | GTAGCTTCGCTGGTAGACATCC |
| Vinculin | GCCAAGCAGTGCACAGATAA | TTCCTTTCTGGTGTGTGAAGC |
| Wnt11 | CTGCACCTCTGGCGACCT | CAGAAGTCAGGGGAGCTCTGT |
| Wnt3 | AGCGTAGCAGAAGGTGTGAAG | GTGGCCCCTTATGATGTGAGT |
| Wnt5a | GTCCTTTGAGATGGGTGGTATC | ACCTCTGGGTTAGGGAGTGTCT |
| Wnt5b | CCCCAGGCCAGAGAAAGC | CCTCCCCGATGTAGGACAT |
